# A Lysate Proteome Engineering Strategy for Enhancing Cell-Free Metabolite Production

**DOI:** 10.1101/2020.04.05.026393

**Authors:** David C. Garcia, Jaime Lorenzo N. Dinglasan, Him Shrestha, Paul E. Abraham, Robert L. Hettich, Mitchel J. Doktycz

## Abstract

Cell-free systems present a significant opportunity to harness the metabolic potential of diverse organisms. Removing the cellular context provides the ability to produce biological products without the need to maintain cell viability and enables metabolic engineers to explore novel chemical transformation systems. Crude extracts maintain much of a cell’s capabilities. However, only limited tools are available for engineering the contents of the extracts used for cell-free systems. Thus, our ability to take full advantage of the potential of crude extracts for cell-free metabolic engineering is limited. Here, we employ Multiplex Automated Genomic Engineering (MAGE) to tag proteins for selective removal from crude extracts so as to specifically direct chemical production. Specific edits to central metabolism are possible without significantly impacting cell growth. Selective removal of pyruvate degrading enzymes are demonstrated that result in engineered crude lysates that are capable of 10 to 20-fold increases of pyruvate production when compared to the non-engineered extract. The described approach melds the tools of systems and synthetic biology to develop cell-free metabolic engineering into a practical platform for both bioprototyping and bioproduction.

**Highlights:**

- A novel method of engineering crude cell lysates for enhancing specific metabolic processes is described.
- Multiplex Automated Genomic Engineering (MAGE) can be used to engineer donor strains for improving cell-free metabolite production with minimal impact on cell-growth.
- The described lysate engineering strategy can specifically direct metabolic flux and create metabolic states not possible in living cells.
- Pooling of the central precursor pyruvate was significantly improved through use of this lysate proteome engineering strategy.

## 1. Introduction

Driven by the prospect of biological systems that can be easily manipulated, the application of synthetic biology tools to *in vitro* environments offers a promising approach to harnessing an organism’s rich metabolic potential^1^. Cell-free systems use cytoplasmic components, devoid of genetic material and membranes, as a means of producing complex chemical transformations. While living cells require membranes, growth substrates, and biochemical regulation, *in vitro* systems sidestep these barriers to manipulation and present an opportunity to explicitly define a system for creating novel proteins and metabolites^2^. In this way, cell free metabolic engineering (CFME) can use the organism’s existing biochemical functions and further combine these capabilities with heterologous pathways to produce chemical precursors, biofuels, and pharmaceuticals.

Efforts to engineer cell free systems have taken different approaches. Ideally, a CFME system would contain a minimal set of components necessary to carry out a desired biochemical process. Previous approaches employed a defined set of purified enzymes for producing high-yielding chemical conversions and have successfully demonstrated a variety of capabilities including chemical production and protein synthesis^3,4^. Constructing complex, multistep pathways require significant development and upfront costs as utilizing purified proteins at scale remains costly^5^. Further, these purified component systems can exhibit slow catalysis rates, possibly due to the lack of accessory proteins and appropriate protein concentrations capable of improving pathway yield^5^. Nevertheless, long running CFME systems that can catalyze multi-step reaction pathways for days have been developed^3^.

The use of crude cell extracts presents an alternative approach to CFME. Simple cell lysis and minimal fractionation can be rapidly carried out and result in complex enzyme mixtures for a fraction of the cost of purified components^6–8^. Crude extract systems derived from both commonly used cell-free model organisms, such as *E. coli* BL21 Star (DE3), or nontraditional strains, such as *Vibrio natriegens*, contain a similar biochemistry to the donor cell and can serve as both prototyping tools for *in vivo* metabolic engineering and as bioproduction platforms^9–12^. Cell-free systems work well for this purpose as extracts can be modularly assembled with those enriched for metabolic pathways in order to produce a specific molecule. Additionally, their compatibility with chemical reactors and ability to consume low cost feedstocks have popularized them as potential sources for industrial production^13–15^.

While environmental variables of a cell-free system can easily be manipulated, the proteomic content of the crude extract is more difficult to engineer. Genetic manipulation of a donor strain can substantially impact its growth and function as a bioproduction system^16^. It has been noted previously that simple variations in growth conditions can lead to complex changes in the proteome and significant differences in metabolite flux in the resulting crude extracts^16,17^. Further, specific enzymes can be added or expressed in an extract to further define metabolite production^1,7^. However, removing specific proteins is challenging as gene deletions can affect the growth and global expression of the donor cell. In particular, deletions to central metabolism can be lethal, which severely limits the ability to direct flux from simple carbon sources. The inability to remove specific pathways from CFME reactions poses a significant constraint and limits the use of crude extracts for bioproduction^7,16^. Tools that allow shaping of the proteome and tailoring of specific pathways will be critical in enabling the use of crude extract systems for metabolic engineering applications.

In this work, we describe the use of genome engineering, specifically MAGE, to enable the removal of particular proteins from crude extracts for CFME. 6xHis-tags are incorporated into proteins that are expected to consume pyruvate and are used for the affinity-based removal of these proteins following cell lysis. Pyruvate, a key connection point in central carbon metabolism, was chosen due to its role linking glycolysis and the Krebs cycle, as well as its relevance as a central precursor for numerous products^18,19^. The use of genome engineering results in a modified proteome capable of producing engineered metabolic phenotypes with minimal impact on the viability of the donor cell (Figure 1)^20,21^. This general strategy was demonstrated using a *E. coli* BL21 Star (DE3) as a chassis due to its prevalent use for cell-free synthetic biology.

**Figure 1.**
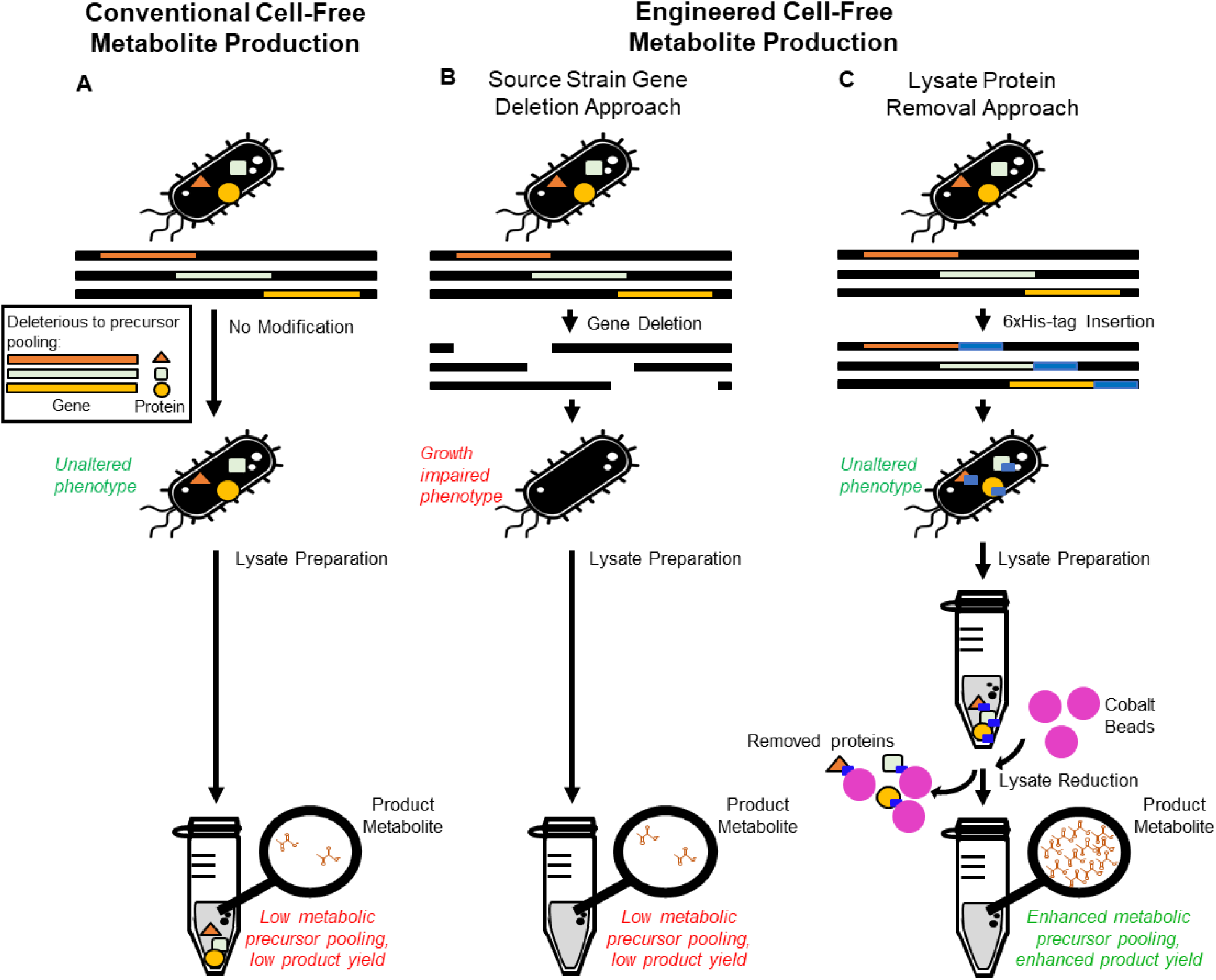
Overview of approaches to preparing lysates for cell free metabolic engineering. **A.** Complex metabolism present in *E. coli* lysates harnessed for cell-free metabolite production can compromise central metabolic precursor yields. **B & C.** Cell-free metabolic engineering approaches seek to reduce lysate complexity in order to redirect carbon flux and pool central metabolic precursors. **B.** The standard CFME approach reduces lysate complexity by deleting target genes from the source strain, often resulting in growth impaired or lethal phenotypes due to the inability to remove essential genes. This can require multiple design-build-test cycles. **C.** The new approach involves engineering source strains to endogenously express recognition sequences, such as 6xHis-tags, into target proteins for subsequent removal from lysates through affinity purification, resulting in minimal to no impact on source strain growth and enhanced pooling of specific metabolic products.

## 2. Methods

### 2.1 Generation and Validation of Genome Engineered Strains using MAGE

All multiplex allele-specific PCR (MASC-PCR), Sanger Sequencing oligos, and recombineering oligos were created manually and ordered from IDT (Coralville, IA) with standard purification. Each targeting oligo incorporated four phosphorothioated bases on the 5′ terminus. An 18-base CACCATCACCATCACCAT sequence was used to add the 6xHis-tag and directed at either the N- or C-terminus based on previous literature or crystal structure analysis^22,23^. The pORTMAGE protocol used in this study followed previous work with the exception that growth was carried out in 6 mL of Luria-Bertani-Lennox (lbl) cultures in glass tubes with 100 mg/mL of carbenicillin, recovery was performed in 3 mL of terrific broth with a 1-hour incubation time prior to adding 3 mL of lbl-carb for outgrowth^24^. Given the significant time required to find accumulated mutations in a single strain, the additive mutations were started from previously found mutations such that Δ1 was used to create Δ2 and so on as per the protocols used in previous studies^4^. After every 8-12 cycles of MAGE, 30-60 colonies were screened for genome edits using MASC-PCR as detailed previously^4^. Allelic genotyping was performed using standard primers designed to flank both modified genes. Amplicons were Sanger sequenced to validate the insertion of the 6xHis-tag sequence. Primer sequences used in this study are listed in Supplemental Table 1 and Supplemental Table 2.

### 2.2 Cell-Free Extract Preparation Protocol

Following plasmid curing, the cell extracts were prepared from *E. coli* BL21 Star (DE3) grown at 37 °C in 2xYPT-G (16 g L-1 tryptone, 10 g L-1 yeast extract, 5 g L-1 NaCl, 7 g L-1 KH2PO4, 3 g L-1 K2HPO4, 18 g L −1 glucose). Cell extracts were prepared by harvesting 50-mL cultures grown in baffled Erlenmeyer flasks to an OD600 of 4.5 Cells were harvested by centrifugation at 5000x g for 10 min in 50 mL volumes and washed twice with S30 buffer (14 mM magnesium acetate, 60 mM potassium glutamate, 1 mM dithiothreitol (DTT) and 10 mM Tris-acetate, pH 8.2) by resuspension and centrifugation. The pellets were weighed, flash-frozen, and stored at –80 °C. Extracts were prepared by thawing and resuspending the cells in 0.8 mL of S30 buffer per mg of cell wet weight. The resuspension was lysed using 530 joules per mL of suspension at 50% tip amplitude with ice water cooling. Following sonication, tubes of cell extract were centrifuged twice at 21,100 x g for 10 minutes at 4 °C, aliquoted, frozen with liquid nitrogen, and stored at –80 °C.

### 2.3 Cell-Free Extract Reductions

Cell extracts were reduced by adding one volume of cell extract to 0.2X volume of ice-cold HisPur™ Cobalt Resin (ThermoFisher Scientific) suspension in 1.5 mL microcentrifuge tubes. Prior to the addition of lysate, HisPur™ Cobalt Resin was washed 2X with 500 µL S30 buffer and incubated with 10 mM imidazole buffer (pH 4.5; 10 mM imidazole, 50 mM monopotassium chloride, 300 mM NaCl). Lysate-resin mixtures were incubated for 1 hour at 4°C under shaking conditions (800 rpm) to ensure the suspension of the resin particles in the extracts and then centrifuged at 14,000 x g for 30 seconds. Supernatants were aliquoted, flash-frozen, and stored at −80°C until used. His-tagged proteins were eluted from the HisPur™ Cobalt Resin by suspending the resin in 50 µL elution buffer (pH 4.5; 250 mM Imidazole, 50 mM monosodium phosphate, 300 mM NaCl) for 30 minutes at 4°C under shaking conditions (800 rpm). The eluate was obtained for proteomic quantification by spinning down the suspension at 14,000 x g for 30 seconds and collecting the supernatant.

### 2.4 CFME Reaction Set-up

Glucose consumption reactions were carried out at 37 °C in 50 μL volumes using a solution of 250 mM glucose, 18 mM magnesium glutamate, 15 mM ammonium glutamate, 195 mM potassium glutamate, 1 mM ATP, 150 mM Bis-Tris, 1 mM NAD+, 10 mM dipotassium phosphate. Similarly, pyruvate fed reactions were carried out using the same conditions with the exception of 25 mM pyruvate being used in place of glucose. Extracts were added to a final protein concentration of 4.5 mg mL-1. Each reaction was quenched by the addition of 50 μL of 5% TCA. The supernatant was centrifuged at 11,000 x g for 5 minutes and directly used for analytical measurements.

### 2.5 Proteomics Sample Preparation

Samples of both reduced and unreduced versions of WT, 6xHis-1, 6xHis-2, 6xHis-3, and 6xHis-4 cell extracts were each prepared in triplicate as follows. Extracts were solubilized in 200 μL of 4% SDS in 100 mM Tris buffer, pH 8.0. Trichloroacetic acid was added to achieve a concentration of 20% (w/v). Samples were vortexed and incubated at 4°C for 2 h followed by 10 min at −80°C. Samples were then thawed on ice prior to centrifugation (∼21,000g) for 10 min at 4°C to pellet precipitated proteins from the detergent and solutes. The supernatant was discarded, and samples were washed with 1 mL of ice-cold acetone. Pelleted proteins were then air-dried and resuspended in 100 μL of 8 M urea in 100 mM Tris buffer, pH 8.0. Proteins were reduced with 10mM dithiothreitol incubated for 30 min and alkylated with 30mM iodoacetamide for 15 min in the dark at room temperature. Proteins were digested with two separate and sequential aliquots of sequencing grade trypsin (Promega) of 1 μg. Samples were diluted to 4 M urea and digested for 3 hours, followed by dilution to 2 M urea for overnight digestion. Samples were then adjusted to 0.1% trifluoracetic acid and then desalted on Pierce peptide desalting spin columns (Thermo Scientific) as per manufacturer’s instructions. Samples were vacuum-dried with a SpeedVac (Thermo Scientific) and then resuspended in 50 μL of 0.1% formic acid. Peptide concentrations were then measured using a NanoDrop spectrophotometer (Thermo Scientific) and 2 μg of each sample was used for LC-MS measurement.

### 2.6 LC-MS/MS analysis

All samples were analyzed on a Q Exactive Plus mass spectrometer (Thermo Scientific) coupled with an automated Vanquish UHPLC system (Thermo Scientific). Peptides were separated on a triphasic precolumn (RP-SCX-RP; 100 μm inner diameter and 15 cm total length) coupled to an in-house-pulled nanospray emitter of 75 μm inner diameter packed with 25 cm of 1.7 μm of Kinetex C18 resin (Phenomenex). For each sample, a single 2 μg injection of peptides were loaded and analyzed across a salt cut of ammonium acetate (500 mM) followed by a 210 min split-flow (300 nL/min) organic gradient, wash, and re-equilibration: 0% to 2% solvent B over 27 min, 2% to 25% solvent B over 148 min, 25% to 50% solvent B over 10 min, 50% to 0% solvent B over 10 min, hold at 0% solvent B for 15 min. MS data were acquired with the Thermo Xcalibur software using the top 10 data-dependent acquisition.

### 2.7 Proteome Database Search

All MS/MS spectra collected were processed in Proteome Discoverer, version 2.3 with MS Amanda and Percolator. Spectral data were searched against the most recent *E. coli* reference proteome database from UniProt to which mutated sequences and common laboratory contaminants were appended. The following parameters were set up in MS Amanda to derive fully tryptic peptides: MS1 tolerance = 5 ppm; MS2 tolerance = 0.02 Da; missed cleavages = 2; Carbamidomethyl (C, + 57.021 Da) as static modification; and oxidation (M, + 15.995 Da) as dynamic modifications. The Percolator false discovery rate threshold was set to 1% at the peptide-spectrum match and peptide levels. FDR-controlled peptides were then quantified according to the chromatographic area-under-the-curve and mapped to their respective proteins. Areas were summed to estimate protein-level abundance.

### 2.8 Proteomic Data Analysis

For differential abundance analysis of proteins, the protein table was exported from Proteome Discoverer. Proteins were filtered to remove stochastic sampling. All proteins present in 2 out of 3 biological replicates in any condition were considered valid for quantitative analysis. Data was log2 transformed, LOESS normalized between the biological replicates and mean-centered across all the conditions. Missing data were imputed by random numbers drawn from a normal distribution (width = 0.3 and downshift = 2.8 using Perseus software (http://www.perseus-framework.org). The resulting matrix was subjected to ANOVA and a post-hoc TukeyHSD test to assess protein abundance differences between the different experimental groups. The statistical analyses were done using an in-house developed R script.

### 2.9 Glucose and pyruvate measurements

Glucose and pyruvate measurements were performed using high-performance liquid chromatography (HPLC). An Agilent 1260 equipped with an Aminex HPX 87-H column (Bio-Rad, Hercules, CA) and a diode array UV-visible detector (Agilent, Santa Clara, CA) reading at 191 nm. Analytes were eluted with isocratic 5 mM sulfuric acid at a flow rate of 0.55 mL min–1 at 35 °C for 25 min. Pyruvate and glucose standards were used for sample quantification using linear interpolation of external standard curves.

## 3. Results and Discussion

### 3.1 Preparation of 6xHis-tagged strains using MAGE

Pyruvate sits at the biochemical junction of glycolysis and the TCA cycle. It is a key intermediate in producing many food, cosmetic, pharmaceutical and agricultural products whose improved production has been largely unexplored in cell-free systems^18,19^. In order to create a pyruvate pooling phenotype in an *E. coli* cell-free extract, four proteins were chosen as targets for removal, LdhA, PpsA AceE, and PflB, due to their direct role in consuming pyruvate (Table 1) (Figure 2)^19^. 6xHis tags were fused on either the amino or carboxyl terminus by genetic modification based on an evaluation of literature related to previous purification attempts or crystal structures in order to find a non-inhibitory location (Table 1)^22,23^.

**Table 1.**
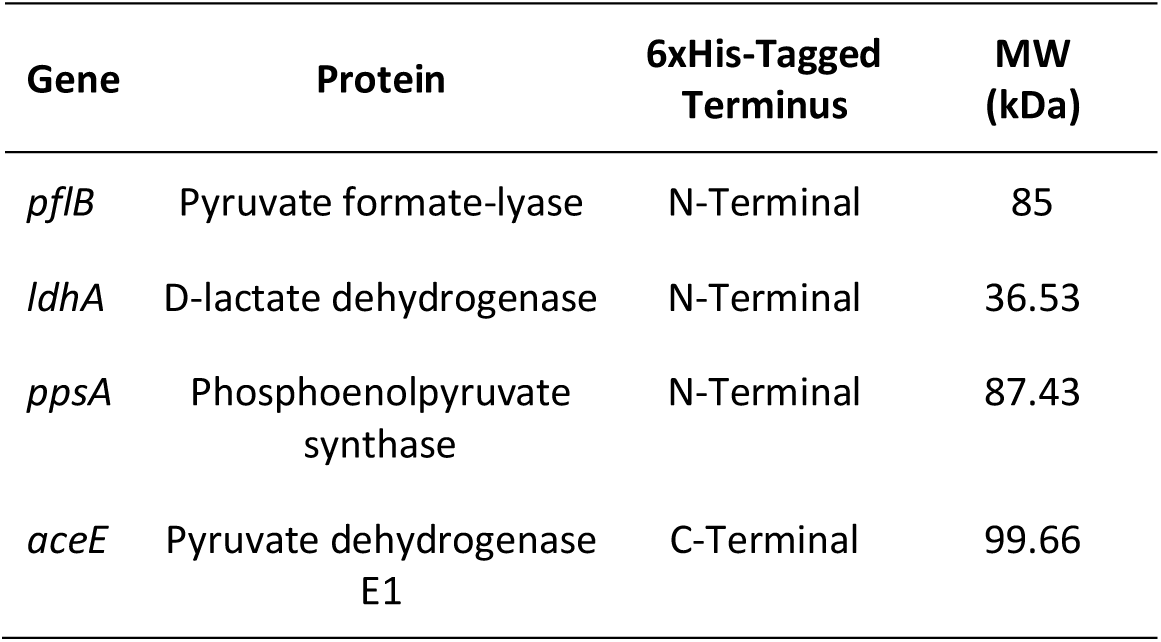
Gene and protein information for MAGE targets with a potential effect on pyruvate metabolism.

| Gene | Protein | 6xHis-Tagged Terminus | MW (kDa) |
| --- | --- | --- | --- |
| *pflB* | Pyruvate formate-lyase | N-Terminal | 85 |
| *ldhA* | D-lactate dehydrogenase | N-Terminal | 36.53 |
| *ppsA* | Phosphoenolpyruvate synthase | N-Terminal | 87.43 |
| *aceE* | Pyruvate dehydrogenase E1 | C-Terminal | 99.66 |

**Figure 2.**
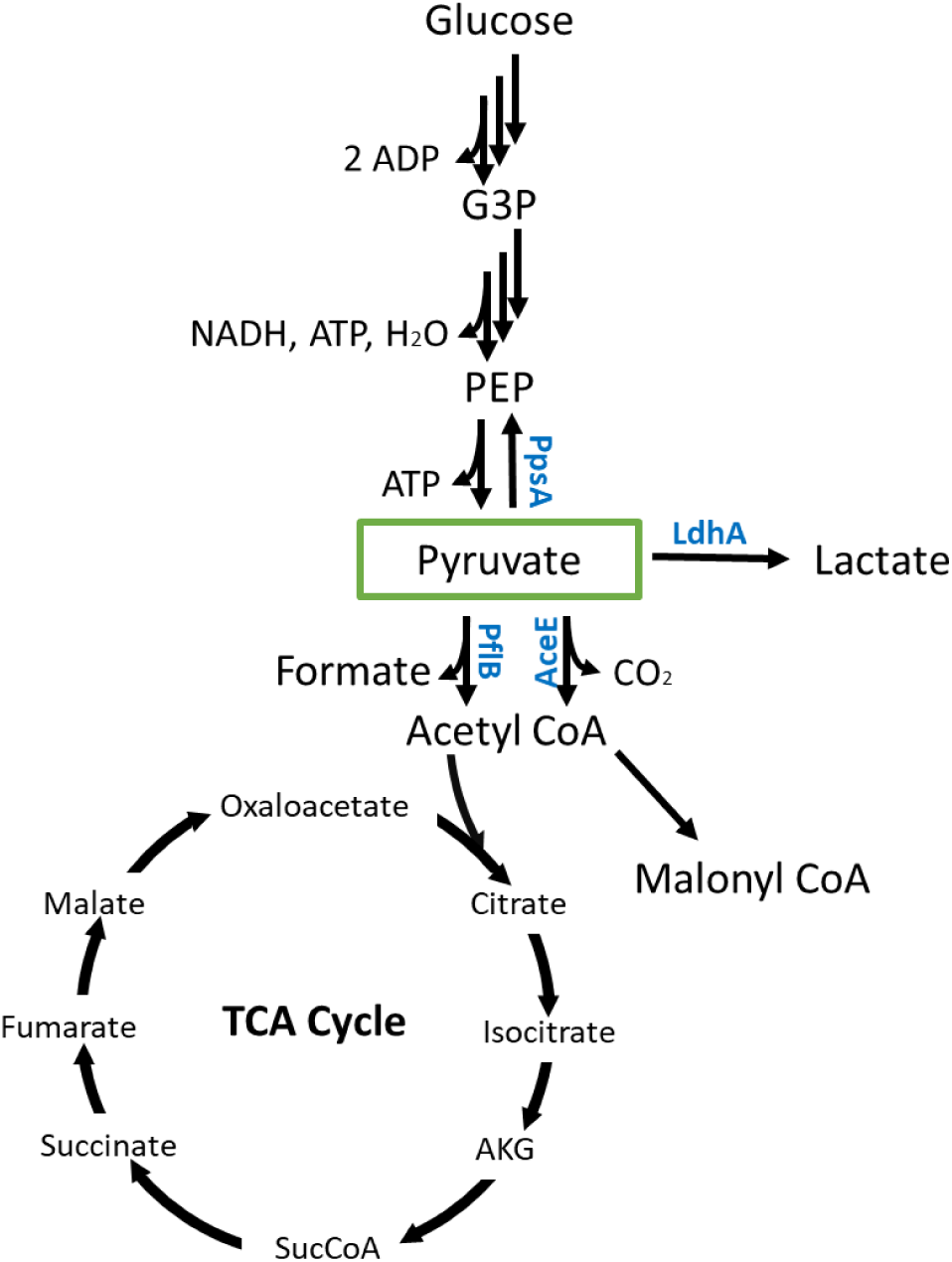
Glycolysis and engineered pathway nodes showing the location of the modified enzymes PflB, LdhA, PpsA, AceE.

The traditional cell-free protein synthesis chassis *E. coli* BL21 Star(DE3) was chosen as the source strain for creating edited extracts. For genome engineering, the pORTMAGE system was used instead of a genome integrated system due to its potential transferability to multiple donor organisms including *E. coli* BL21 Star(DE3) ^25,26^. Additionally, pORTMAGE is curable following genome engineering and relieves the metabolic burden on the cell that can be imparted due to plasmid maintenance^27^. Colony screening was performed using MASC-PCR and further verified using Sanger sequencing (Figure 3 AB)^28^. A total of 5 strains were used for this work. (Table 2). The strains included 6xHis-1 (6xHis-*pflB*), 6xHis-2 (6xHis-*pflB-ldhA*), 6xHis-3 (6xHis-*pflB-ldhA-ppsA*), 6xHis-4 (6xHis-*pflB-ldhA-ppsA-aceE*) and 6xHis-*ldhA*. 60 rounds of MAGE were needed to incorporate all four of the tags into the *E. coli* genome (Figure 3 A) (Table 2). This is high when compared to studies producing single nucleotide changes, but in line with other efforts using 6xHis-tags with a genome incorporated MAGE system^4^.

**Table 2.** Strains created for this study.

| Strain Name | Background | Modified Genes |
| --- | --- | --- |
| 6xHis-ldh | BL21 Star(DE3) | 6xhis-ldhA |
| 6xHis-1 | BL21 Star(DE3) | 6xhis-pflB |
| 6xHis-2 | BL21 Star(DE3) | 6xhis-ldhA, 6xhis-pflB |
| 6xHis-3 | BL21 Star(DE3) | 6xhis-ldhA, 6xhis-pflB, 6xhis-ppsA |
| 6xHis-4 | BL21 Star(DE3) | 6xhis-ldhA, 6xhis-pflB, 6xhis-ppsA, 6xhis-aceE |

**Figure 3.**
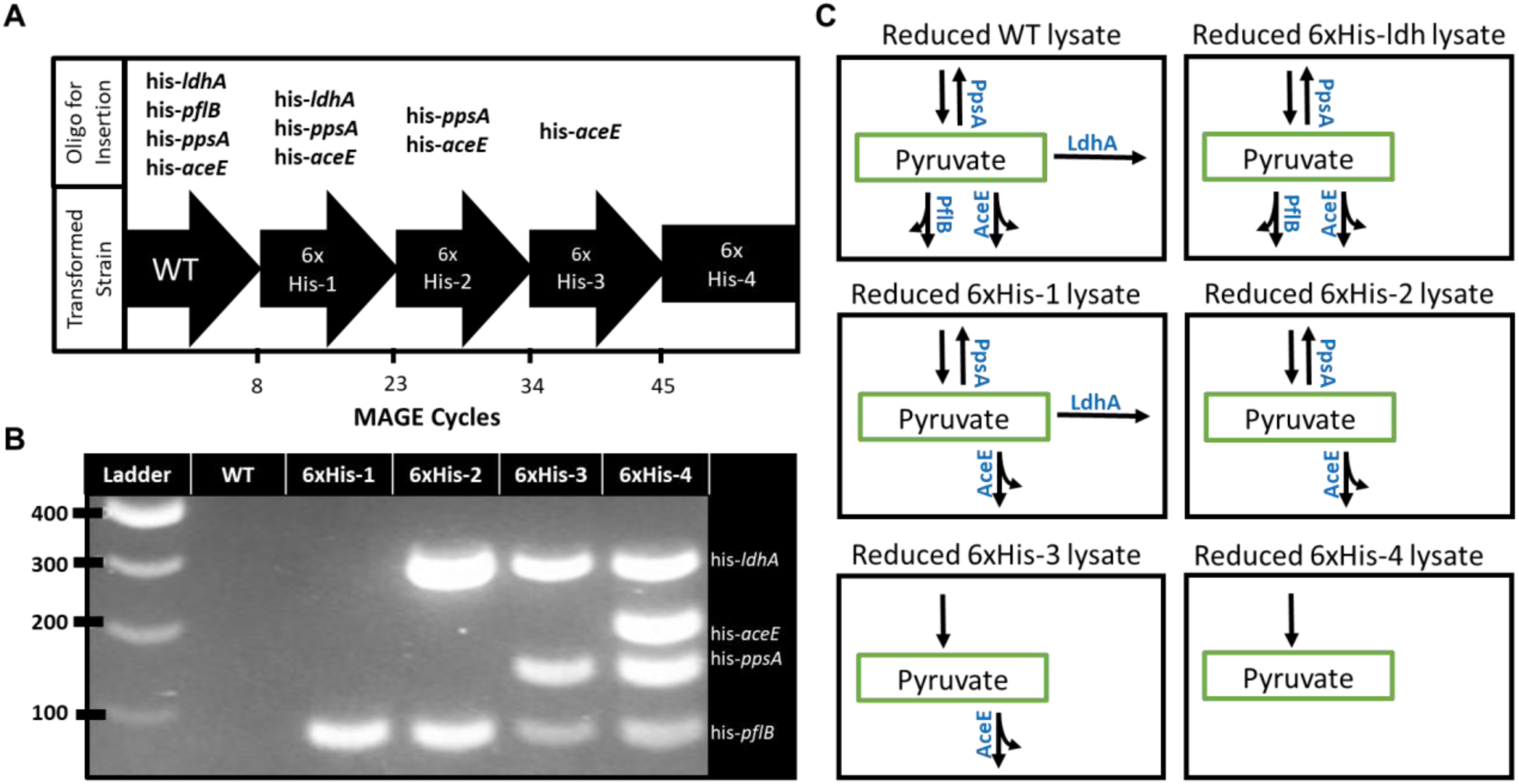
Source strain multiplex genome engineering and expected metabolic phenotypes of derived lysates post-reduction. **A.** Strain construction course by MAGE cycling culminating with the 6xHis-4 containing all 4 tags. Each arrow designates the strain being taken through the MAGE process with the oligos used to transform each strain above the arrow. **B.** MASC-PCR results for additive mutations using primers specifically designed for the 6xHis-taggged version of the gene. **C.** Expected metabolic phenotypes present in WT and engineered lysate proteomes after the reduction of lysates derived from all generated strains.

After curing each strain of the pORTMAGE plasmid, potential inhibitory effects on growth caused by the expression of tagged proteins were evaluated. Though the presence of the polyhistidine tags has previously been observed to cause growth defects due to the stability of tagged proteins, none of the cells produced for this work showed a significant drop in growth rate (Supplemental Figure 1)^29,30^.

### 3.2 In vitro production and measurement of pyruvate

The effect of proteome reduction on the extract’s metabolic profile was then tested by measurement of pyruvate accumulation in a CFME reaction mix. Early time points show high accumulation of pyruvate starting at 2 hours and continuing up to 12 hours before the concentration of the metabolite decreases (Figure 4). After 8 hours, the reduced 6xHis-4 extract reaches a maximum concentration of 25 mM pyruvate from an initial glucose concentration of 100 mM (Figure 4A and B). The accumulation of pyruvate at early time points is consistent with a reduction in the proteins capable of consuming pyruvate. In general, the extracts reduced for ldhA show significant increases in pyruvate pooling potentially indicating it is the main *in vitro* consumer of pyruvate. Interestingly, the effect was not permanent as the pyruvate concentration decreased in the reduced extracts after 12 hours, falling to 6.41 mM for 6xHis-2, 7.2 mM for 6xHis-3, and complete consumption for 6xHis-4 by 24 h (Figure 4). For the unreduced strains, extracts not incubated with cobalt beads, little to no pooling of pyruvate occurred from 0 to 8 hours; a result that is corroborated by previous observations of *E. coli* extracts^16^. This is notable as the highest pyruvate concentration between 0 to 8 hours, 4.15 mM in unreduced 6xHis-3, was 5-fold lower than reduced versions of 6xHis-2, 6xHis-3, and 6xHis-4 (Figure 4).

**Figure 4.**
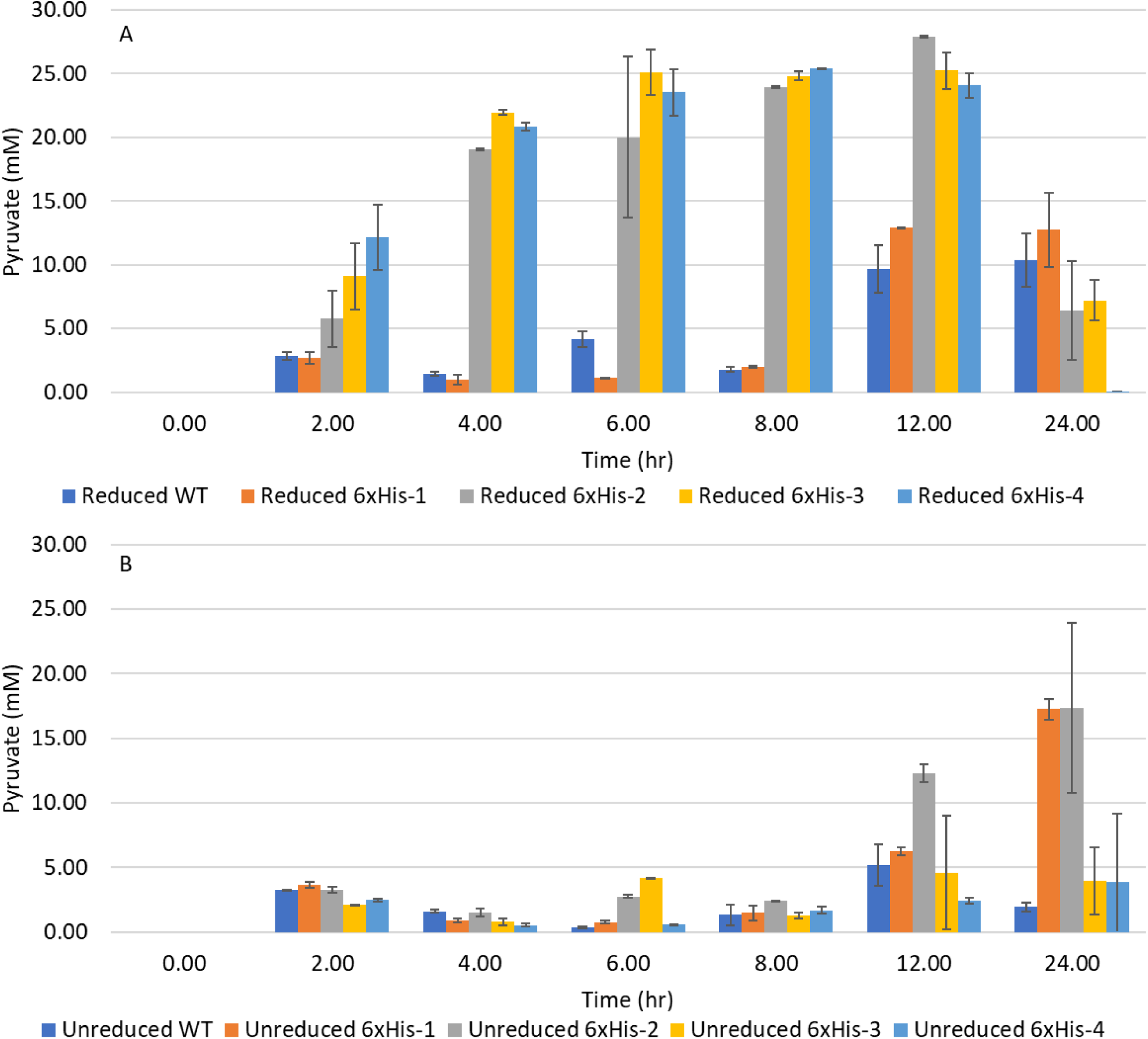
Comparisons of pyruvate concentration over time in WT, 6xHis-1, 6xHis-2, 6xHis-3, and 6xHis-4 lysates. Data and standard deviation for the time course reactions were acquired using n=3 biological replicates. **A.** Extracts that have been reduced for specific proteins 6xHis-tagged proteins. **B.** Extracts without any peptides removed from the proteome.

The rapid accumulation of pyruvate in the reduced strains after 4 hours leads to a relatively stable concentration in reduced 6xHis-2, 6xHis-3, and 6xHis-4 extracts showing that the reduced lysates are able to maintain an equilibrium between glucose and pyruvate concentrations (Figure 4, Supplemental Figure 2). The rapid consumption of pyruvate following 12 hours for the 6xHis-2, 6xHis-3, and 6xHis-4 extracts may indicate that residual pyruvate consuming machinery maintains activity, but enzymes responsible for feeding PEP or cofactors into the system begin to degrade^31^. Additionally, while the concentration of NAD^+^ is bolstered by its exogenous addition during the reaction setup, the elimination of an NADH sink, LdhA, may lead to decreased pooling of pyruvate at later timepoints^7^. These time-dependent changes may indicate that the glycolytic machinery deteriorates faster than the remaining pyruvate degrading pathways. Alternatively, the rapid consumption of the NAD^+^ supply could be limiting due to the potential lack of cofactor recycling initiated by the removal of ldhA. Pyruvate consumption experiments performed with the WT and 6xHis-4 lysates show that a significant portion of the pyruvate consuming pathways remain robust after reduction evidencing that the constraint may be due to bottlenecks in upstream glycolysis (Supplemental Figure 3).

### 3.3 Analysis of LdhA Reduction

To validate the observed additive effects of successful LdhA and PpsA removals on metabolic output, we generated a strain with a single 6xHis-*ldhA* insertion and compared its extent of pyruvate pooling with the 6xHis-4-reduced lysates (Supplemental Figure 4). Importantly, the extract showed that while the reduction of LdhA significantly impacts pyruvate accumulation, it was not solely responsible for the effect. At its peak the 6xHis*-ldhA* reduced extract had only 15.73 mM pyruvate following reduction, falling significantly short of the 27.92 mM, 25.24 mM, and 25.39 mM pyruvate values for 6xHis-2, 6xHis-3, 6xHis-4 mutants, respectively (Figure 3 and Supplemental Figure 4). This result highlights the importance of additive mutations as well as the potential need for developing cell-free specific flux balance models for target selection.

### 3.4 Proteomic of analysis of crude cell extracts

To further understand the pyruvate pooling trends resulting from the use of engineered lysates, differences between the unreduced and reduced proteomes of each of the lysates were evaluated by mass spectrometry (Figure 5). As expected for the WT extract, the reduction process did not significantly impact the presence of the targeted proteins when comparing the peptide abundance values of unreduced and reduced versions (Figure 5A). On the other hand, a substantial 3-to 4-fold decrease in relative LdhA abundance was evident in reduced 6xHis-2, 6xHis-3, and 6xHis-4 lysates compared to their unreduced counterparts (Figure 5C, D, and E), which is consistent with the significant increase in these extracts’ ability to accumulate pyruvate (Figure 3). PpsA was also significantly pulled-down from only the 6xHis-3 and 6xHis-4 lysates. Comparing the metabolic output of these lysates to that of the WT, we observed a 5-to 10-fold increase in pyruvate accumulation at earlier timepoints (Figure 4).

**Figure 5.**
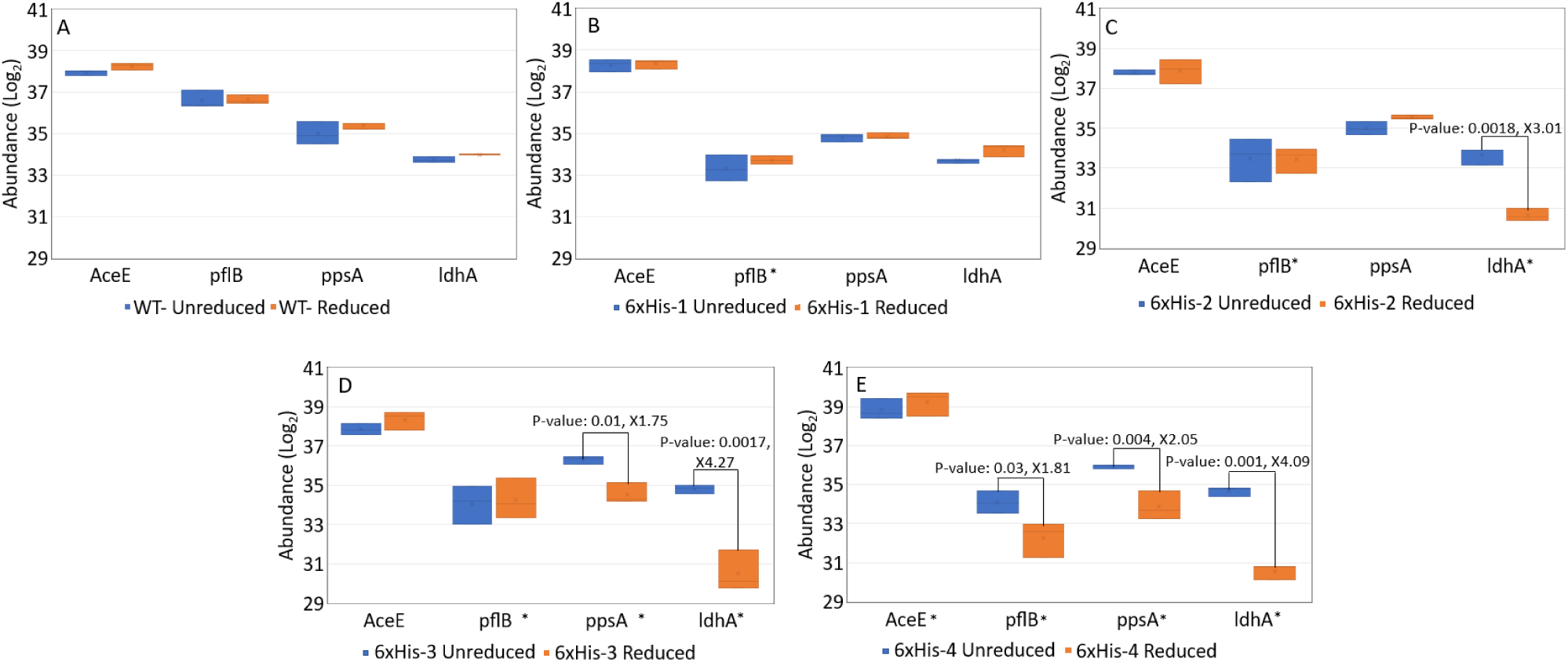
Proteomic analysis of an unreduced cell-free extract (blue), and a reduced cell-free extract (orange). Significant fold-changes in protein concentration when comparing the reduced to the unreduced extract are denoted by p-value and fold change reduction in concentration of the protein above a bracket. Panels A, B, C, D, and E denote the WT, 6xHis-1, 6xHis-2, 6xHis-3, and D. 6xHis-4 strains, respectively. Asterisks indicate proteins targeted for removal in the reduced strain.

From the mass spectrometry-based proteomics profiling, it is evident that 6xHis-tagged LdhA and PpsA could be removed from lysates, while significant removal of 6xHis-tagged PflB was not successful (Figure 5B to E). Although the decrease in PflB levels between the unreduced and reduced 6xHis-4 lysates met the significance threshold (pval < 0.05), the change was only a 1.81-fold reduction compared to the significant decreases in LdhA and PpsA following lysate reduction (Figure 5E). AceE was not observed to be pulled down after the purification of 6xHis-tagged proteins from the 6xHis-4 lysate. These findings are consistent with metabolic output data in that the reduced 6xHis-1 and 6xHis-4 lysates seem to pool as much pyruvate as the reduced WT and 6xHis-3 lysates, respectively (Figure 4).

In their native context, such as in crude cell extracts, both PflB and AceE are known to interact with other proteins. PflB binds to the cytoplasmic membrane protein formate channel A, FocA, to facilitate the translocation of formate and this strong association in *E. coli* lysates has been demonstrated by co-purification experiments^32^. AceE is the pyruvate-binding protein of the pyruvate dehydrogenase complex (PDHC), a multi-enzyme complex that is also comprised of two other components, which is also stable in *E. coli* cell extracts^33^. Individually, overexpressed PflB and AceE have been successfully isolated from their interaction partners through 6xHis-tag purification in the past. However, under native expression and structural context, the use of 6xHis-tags may not be effective for purifying complexed proteins as shorter tags may not destabilize complexes^22,32^. We reason that the formation of these protein-protein interactions may be restricting the accessibility of the target proteins’ termini to the affinity resin, and therefore a larger protein tag that positions itself outside of a complex may be a more favorable alternative^34^. Alternatively, the 6xHis-tags of PflB and AceE are efficiently exposed to the resin, but binding is sterically hindered by the large protein complexes. In this case, the application of protein tags known to efficiently purify intact protein complexes may improve the reduction method^35^.

We further analyzed proteomic changes in the unreduced and reduced extracts to determine whether the reduction method resulted in the removal of off-target proteins in reduced lysates. Importantly, the process of reducing the proteome did not seem to significantly impact proteins in central metabolism outside of those targeted by the reduction process. When comparing the reduced and unreduced WT lysates, in the 58 proteins with a greater than 4-fold reduction, none were connected to central metabolism outside of roles in membrane transport (Supplemental Data 1). Future efforts will seek to minimize off-target effects in order to improve the general applicability of lysate proteome engineering.

Targeted reduction of a lysate proteome enables a rapid means to manipulate central metabolism without the possible drawback of cultivating “sick” organisms as often results from traditional, in vivo metabolic engineering efforts. More broadly, this new approach to engineering in vitro metabolism yields metabolic states that are not traditionally possible in living organisms and sets important groundwork for further developments in cell-free metabolic engineering technology. The pORTMAGE system offers the potential for extension of this engineering strategy to other, non-model cell-free chassis. Future improvements to lysate proteome engineering could also entail the use of multiplex genome engineering methods that are amenable to the insertion of larger tags^26,36,37^. To further advance the reduction of the lysate’s proteome, orthogonal protein degradation systems could be employed wherein proteins are genomically tagged and degraded in a cell-free extract using an exogenous protease. The *mf-lon* protease system serves this function. This protease targets a 27 amino acid long peptide and may overcome issues associated with targeting protein complexes^38,39^.

## 4. Conclusions

Shown in this work was the use of genome engineering to effect *in vivo* protein modifications that lead to directed metabolic activity in the derived lysates. This novel cell-free metabolic engineering strategy allows targeted removal of enzymes that enables focused production of metabolites from simple precursors using rapidly prepared crude extracts that would otherwise lead to changes in metabolic state and significant growth defects in living cells^40–43^. The ability to extract pyruvate degrading enzymes, leading to unconventional metabolic states, was engineered and shown to be capable of pooling pyruvate for a significant period of time. The ability to direct metabolic flux in cell-free systems and create proteomes untenable to living cells was demonstrated. The flexibility of CFME systems highlights the significant value they hold as novel bioproduction platforms. The advances made in this work can be extended to design molecule specific donor strains for natural product biosynthesis, such as for polyketides or carbohydrates, through the removal of defined inhibitory reactions. The ability to reduce specific components of crude lysates allows for more complex reaction networks to be employed in the development of CFME bioproduction platforms. As CFME begins to tackle new challenges related to antibiotic, fuel, and materials production, innovative engineering tools and techniques will be crucial to advancing the scope and adoption of cell-free biological production.

## Contributions

Conceptualization, D.C.G., J.L.D., M.J.D.; Methodology, D.C.G., J.L.D., H.S., P.E.A., R.L.H., M.J.D.; Investigation, D.C.G., J.L.D.; Writing – Review and Editing, All authors.; Resources and supervision, M.J.D., P.E.A., R.L.H.

## Acknowledgment

Special thanks to Ákos Nyerges for providing valuable advice regarding the use of the pORTMAGE system. This research was sponsored by the Genomic Science Program, U.S. Department of Energy, Office of Science, Biological and Environmental Research, as part of the Plant Microbe Interfaces Scientific Focus Area (http://pmi.ornl.gov). Oak Ridge National Laboratory is managed by UT-Battelle, LLC, for the U.S. Department of Energy under contract DE-AC05-00OR22725. D.C.G. is an NSF Graduate Fellow. This manuscript has been authored by UT-Battelle, LLC under Contract DE-AC05-00OR22725 with the U.S. Department of Energy. The United States Government retains and the publisher, by accepting the article for publication, acknowledges that the United States Government retains a non-exclusive, paid-up, irrevocable, worldwide license to publish or reproduce the published form of this manuscript, or allow others to do so, for United States Government purposes. The Department of Energy will provide public access to these results of federally sponsored research in accordance with the DOE Public Access Plan (http://energy.gov/downloads/doe-public-access-plan).

## Appendix A. Supplementary data References

## Supplemental Material

**Supplemental Table 2.**
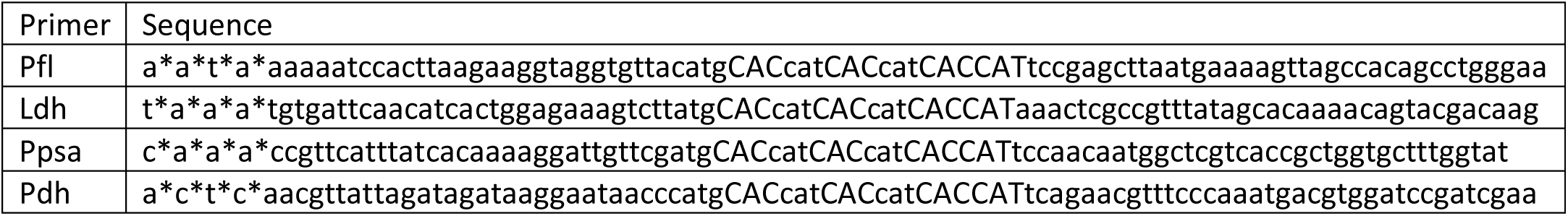
MAGE oligos use for this study. Phosphorothioated bases are noted with asterisks.

**Supplemental Table 3.**
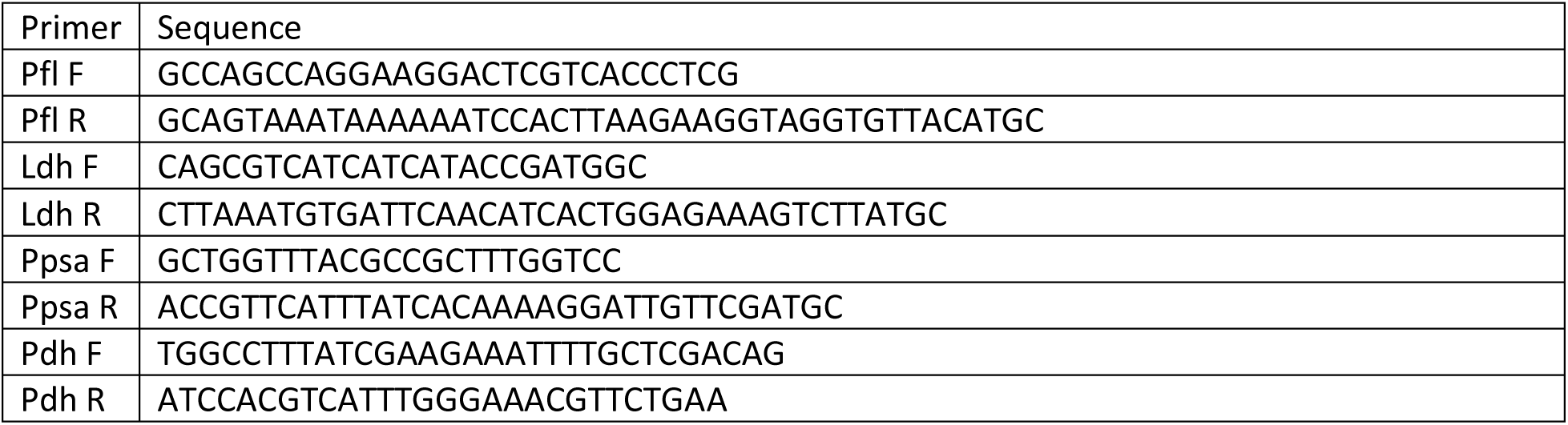
MASC-PCR oligos used for this study.

**Supplemental Figure 1.**
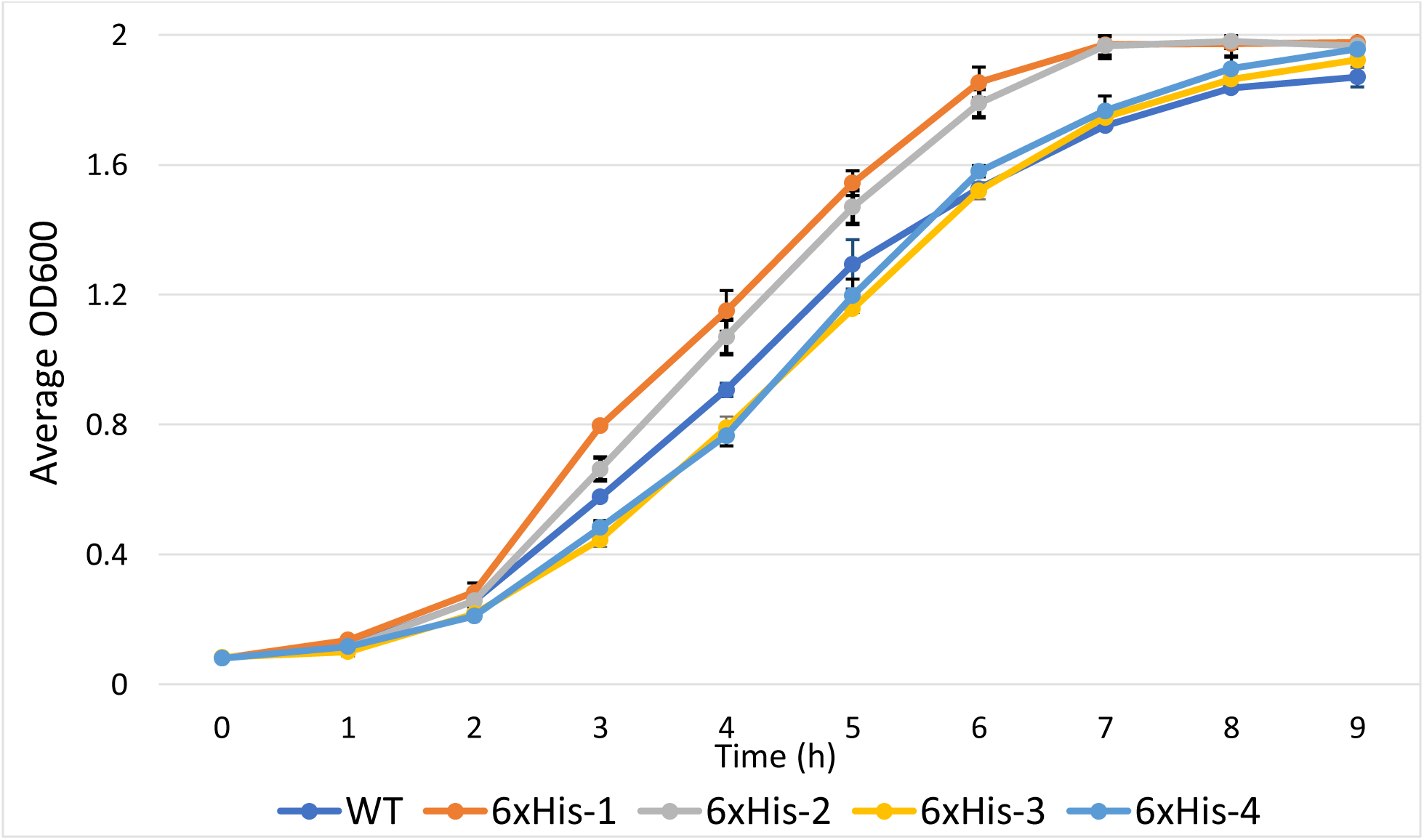
Growth rate and terminal OD600 was measured using for the four main strains produced for this study.

**Supplemental Figure 2.**
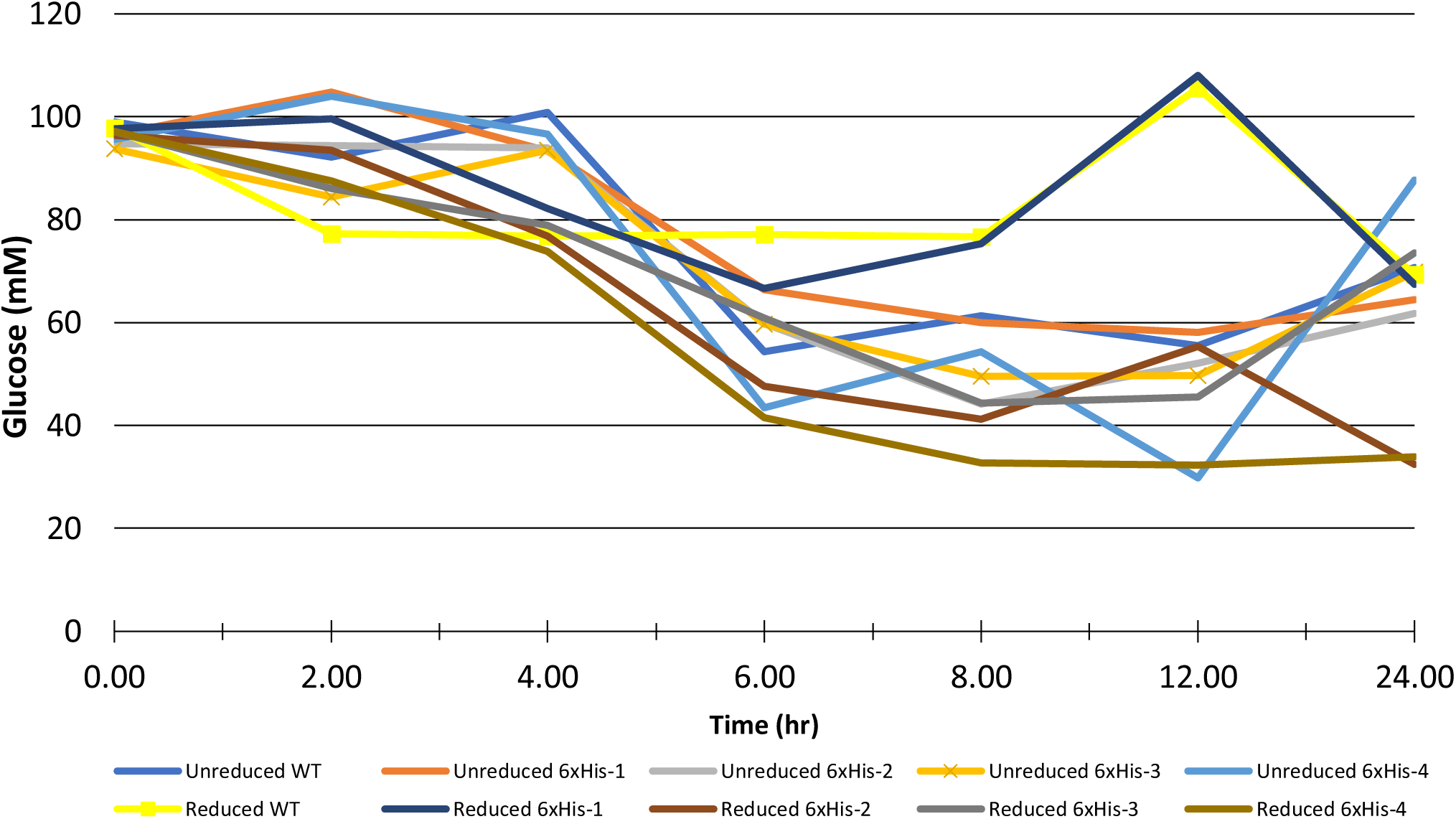
Glucose measured over a 24-hour time period.

**Supplemental Figure 3.**
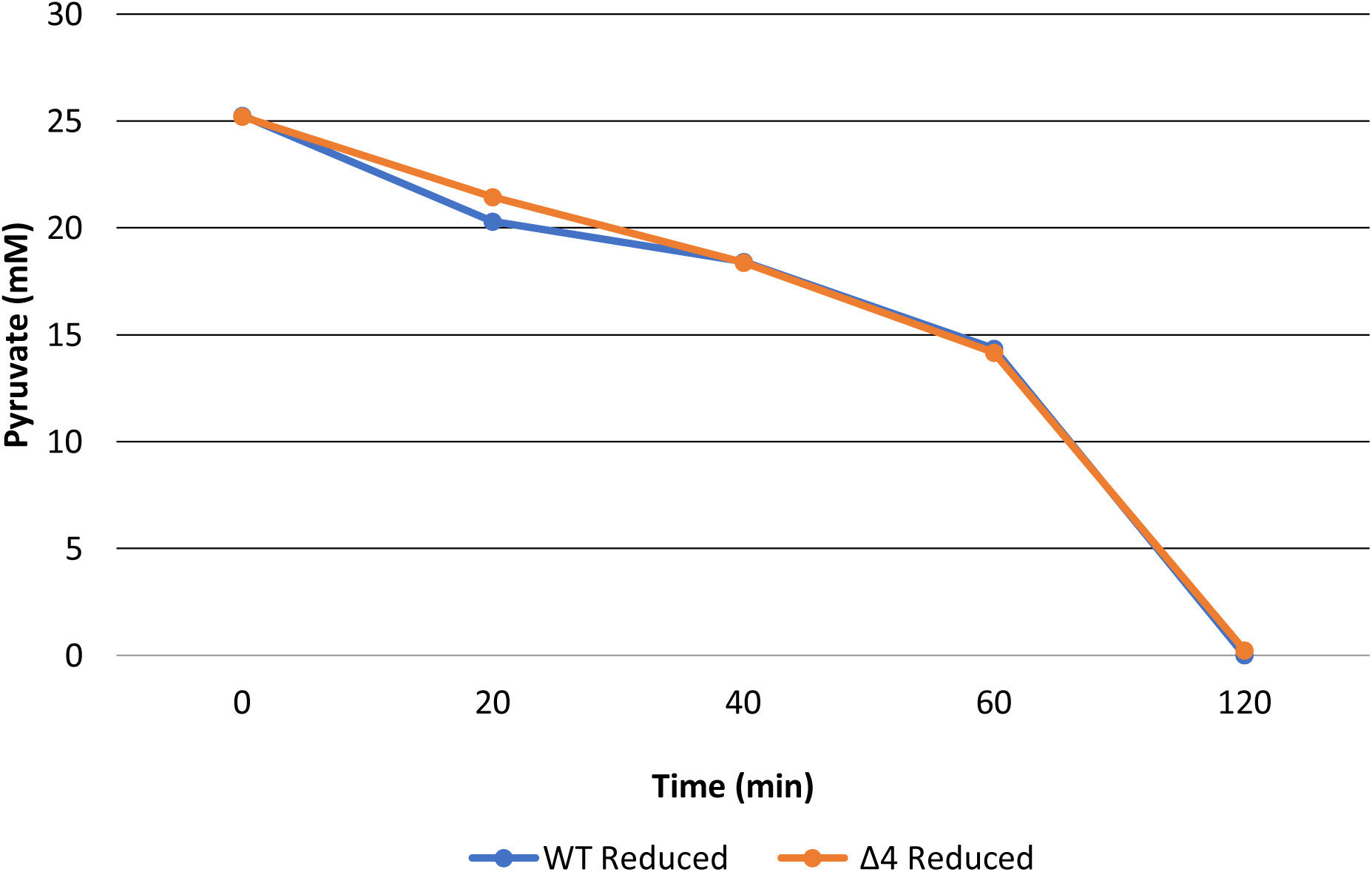
Comparison of pyruvate consumption using reduced WT and Δ4 extracts. All reaction conditions were exactly as those used for the glucose consuming reactions aside from the replacement of glucose with 25 mM pyruvate.

**Supplemental Figure 4.**
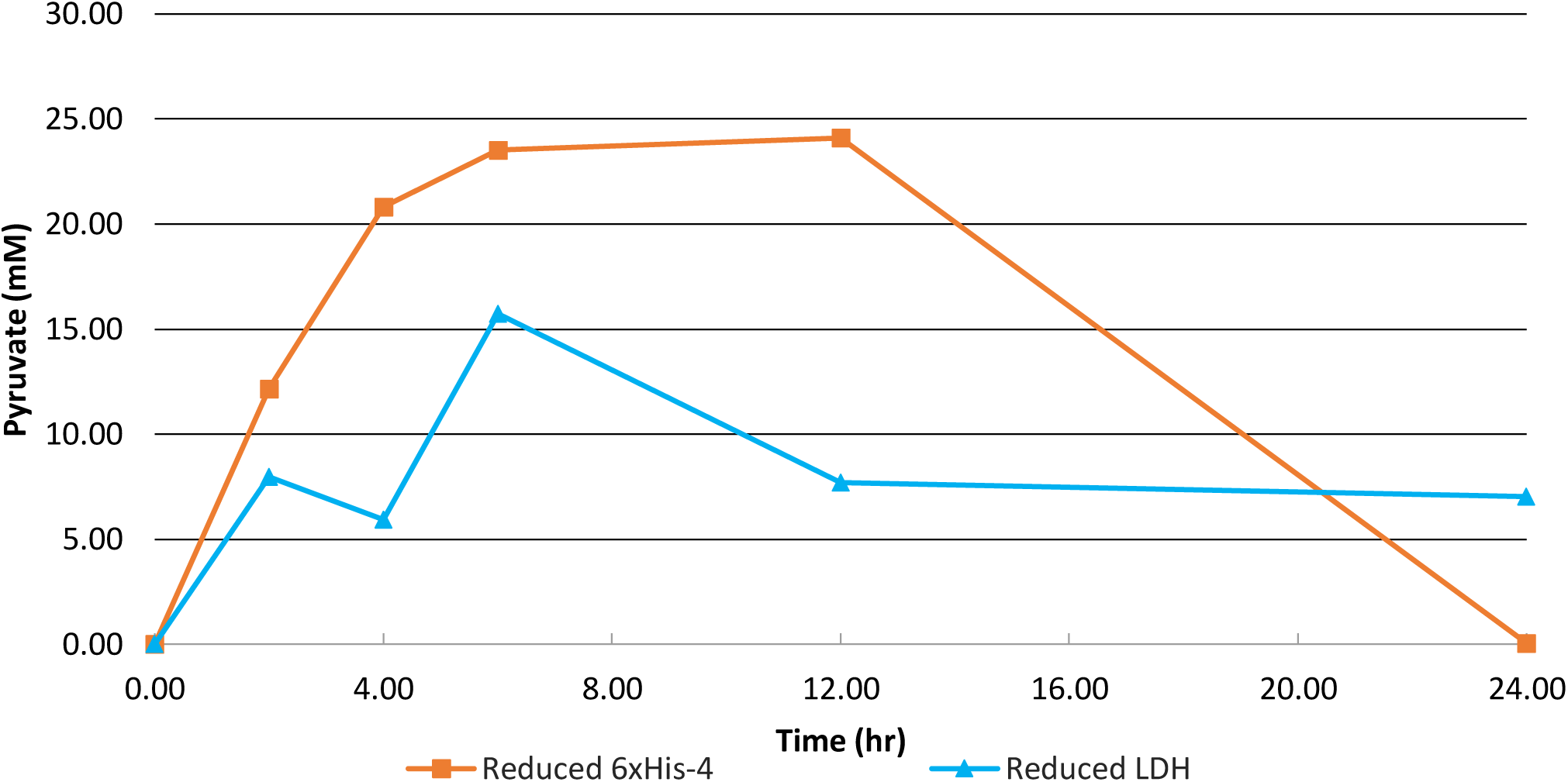
Comparison of pyruvate concentration changes when fed 100 mM glucose for an extract with a single reduction of ldhA to an extract reduced for all 4 proteins his-ldhA, his-pflB, his-ppsa, his-aceE.

